# The Neurodynamic Core of Meditation: Dissociating Meditation from Rest and Task in a Reliability-based EEG study

**DOI:** 10.64898/2026.05.27.728082

**Authors:** Praerna Chowdhury, Ramajayam Govindaraj, Arun Sasidharan, Apar A Saoji, Ravindra P N, Georg Northoff, Bindu M Kutty

**Affiliations:** Centre for Consciousness Studies, Department of Neurophysiology, NIMHANS, Bangalore, Karnataka, INDIA; Indian Knowledge system & Mental Health Applications Centre (IKSMHA), IIT-Mandi, Himachal Pradesh, INDIA; Swami Vivekananda Yoga Anusandhana Samsthana, Bangalore, Karnataka, India; The Royal’s Institute of Mental Health Research & University of Ottawa, Brain and Mind Research Institute, Centre for Neural Dynamics, Faculty of Medicine, University of Ottawa, 145 Carling Avenue, Rm. 6435, Ottawa, ON K1Z 7K4, Canada

**Keywords:** reproducibility, repeatability, neural marker of meditation, multi-day design, meditation proficiency

## Abstract

**Background:** Electroencephalographic (EEG) studies attempting to characterise the neural signature of meditation typically rely on contrasts with passive rest or comparisons among practitioners based on experience. However, these approaches rarely include active control states and seldom establish the reliability and robustness of identified quantitative EEG features. Consequently, the validity of proposed neurophysiological markers of meditative state remains uncertain. The present study addressed these limitations by using a reliability-informed, multi-session within-subject design to characterise distinct state-dependent EEG dynamics in experienced meditators from the Brahmakumaris Rajayoga tradition.

**Methods:** Thirty long-term meditators underwent repeated EEG recordings over two days, comprising two meditation sessions per day. Each meditation block was flanked by rest periods, with a cognitive task between sessions to reduce carryover effects. We quantified broadband spectral power, aperiodic slope and intercept, and nonlinear dynamical measures, including detrended fluctuation analysis (DFA), Higuchi fractal dimension, and permutation entropy (PE), across meditation, rest, and task conditions.

**Results:** Compared with both rest and task states, meditation was associated with increased theta–alpha power, an elevated aperiodic intercept, and systematic modulation of nonlinear indices (DFA, Higuchi, PE). Further meditative core features demonstrated high inter-session test-retest reliability, strong inter-individual consistency, stability across guided and silent meditation states, and were not moderated by years of meditative experience.

**Conclusion:** The present framework identifies a reproducible neurodynamic core of meditation, distinct from passive and active control states, spanning spectral, aperiodic, and nonlinear EEG domains in long-term meditators. These findings enhance the construct validity and measurement reliability of meditation-specific neural markers.

## 1. Introduction

Over the past two decades, neuroscientific investigations of meditative states have expanded substantially. Accumulating evidence suggests that meditation is neuro-physiologically distinct from sleep, drowsiness, focused attention, and other higher-order cognitive states (Hinterberger et al., 2014), particularly in long-term practitioners (Fell et al., 2010). Yet despite these advances, findings remain highly variable and often inconsistent (Lee et al., 2018; Lomas et al., 2015), complicating efforts to establish a stable neural characterisation of meditative state (Brandmeyer et al., 2019; Travis, 2020).

A central challenge lies in defining and operationalising “meditation” as a neural construct. Meditation is not a unitary cognitive process but a heterogeneous family of practices encompassing distinct attentional modes (Travis & Shear, 2010), phenomenological experiences (Travis, 2020), and intended outcomes (Brandmeyer et al., 2019). Focused attention practices (e.g., breath-based (Bréchet et al., 2021), or Samatha meditation), open monitoring approaches (e.g., Vipassana) (Lutz et al., 2008; Pascarella et al., 2025), transcendence-oriented techniques (e.g., Transcendental Meditation) (Josipovic, 2010), devotional absorption (Yordanova et al., 2020), and non-dual awareness traditions (Travis & Shear, 2010) differ not only in technique but also in their underlying goals, such as concentration, insight, emotional regulation, or self-transcendence (Y.-Y. Tang et al., 2015; Travis, 2020). Collapsing such diverse practices under a single umbrella term risks obscuring practice-specific, stable, and reproducible neural mechanisms (Awasthi, 2013), thereby contributing to the inconsistencies across the literature (Lee et al., 2018; Lomas et al., 2015).

Although many studies have adopted neurophenomenological frameworks integrating first-person reports with neural measures (thick versus thin descriptions) (Lutz et al., 2025; Northoff & Ventura, 2025), several foundational methodological issues remain underexamined. These include experimental design, analytical approaches (Gupta & Dhawan, 2022), feature selection (Deolindo et al., 2020), comparator conditions (Davidson & Kaszniak, 2015), intra- and inter-individual variability (R. Tang & Braver, 2020), and operational definition of expertise (Sezer & Sacchet, 2025). Consequently, systematic assessments of methodological validity and reliability are rare, limiting confidence in proposed neural markers of meditation.

Electrophysiological findings further illustrate this variability. Meditation-related changes have been reported across oscillatory power (Lee et al., 2018; Lomas et al., 2015), aperiodic activity (McQueen et al., 2024; Pascarella et al., 2025; Vyšata et al., 2014) and signal complexity (Atad et al., 2025; Deolindo et al., 2020). In oscillatory analyses, increases in lower alpha and upper theta power are frequently described (6–8 Hz) (Aftanas & Golocheikine, 2001; Cahn & Polich, 2006; Lee et al., 2018), yet findings in delta, beta, or gamma bands remain inconsistent (Lee et al., 2018; Lomas et al., 2015). Reports of neural complexity and critical dynamics are contradictory (Atad et al., 2025), and aperiodic activity remains comparatively underexplored (Dziego et al., 2024; McQueen et al., 2024; Pascarella et al., 2025). Moreover, different EEG metrics index fundamentally distinct neural properties, making cross-study comparison difficult. The inconsistency across spectral, aperiodic, and nonlinear measures raises a fundamental question: whether a neurodynamic core of the meditative state exists, independent of methodological and reliability issues.

Several methodological factors likely contribute to the heterogeneity of findings. There is a variety of methodological influencing factors. Most studies compare meditation with passive rest (Cahn et al., 2010; Sharma et al., 2018, 2023), contrast experienced practitioners with novices (Baijal & Srinivasan, 2010; Braboszcz et al., 2017; Katyal & Goldin, 2021), or examine differences across experience levels (Kakumanu et al., 2018; Nair et al., 2017). Active control states are rarely included (DeLosAngeles et al., 2016; Malipeddi et al., 2024; Thomas et al., 2014), and the stability or reliability of identified EEG features is seldom quantified. Variations in instructional format, task demands, and session structure further influence neural outcomes (Atad et al., 2025; Brandmeyer & Delorme, 2013). Trait-level adaptations in long-term practitioners may interact with state-level practice effects (Kakumanu et al., 2019; Nair et al., 2017), reducing within- and between-subject reliability. Moreover, years of practice are frequently used as a proxy for expertise (Brefczynski-Lewis et al., 2007) despite emerging evidence that depth, intensity (Kakumanu et al., 2018) and alignment with a tradition’s philosophical framework may be equally critical (Nair et al., 2018). Collectively, these factors may explain inconsistent or even contradictory findings, including reports of both increased (Atad et al., 2025; Walter & Hinterberger, 2022) and decreased neural complexity (Aftanas & Golocheikine, 2002; Atad et al., 2025; Kakumanu et al., 2018), as well as opposing changes in specific frequency bands (Lee et al., 2018; Lomas et al., 2015). Small sample sizes and single-session designs further compound variability, undermining the validity of inferences regarding meditation-specific neural mechanisms (Kaur & Singh, 2015). These factors are rarely quantified or controlled, undermining the validity of inferences about meditation-specific neural mechanisms.

Quantitative EEG (qEEG), encompassing broadband spectral power, aperiodic, and nonlinear measures, has demonstrated strong reliability across rest (Duan et al., 2021), task (Ding et al., 2022) and mental or cognitive performance paradigms (Li et al., 2024; McEvoy et al., 2000), in both healthy and clinical populations across all ages (Li et al., 2024; Lopez et al., 2023; Uudeberg et al., 2025). However, its reliability during intentionally induced meditative states remains largely unexplored, despite evidence that mental states can substantially modulate EEG dynamics (Young et al., 2021). Establishing the stability of qEEG features across repeated meditation sessions may therefore identify robust neurophysiological markers of meditative state and provide objective reference indices for tracking contemplative skill acquisition.

Given the diversity of contemplative traditions, developing a rigorous methodological framework within a single, well-defined practice offers a critical starting point. We therefore address the methodological and reliability challenges inherent in characterising meditation as a dynamic neural state by studying long-term practitioners of Brahmakumaris Rajayoga meditation (BKRY), a globally practised open-eye meditation tradition taught by the Prajapita Brahmakumaris. BKRY has been classified as a complex meditation method engaging affective, cognitive, and null states during practice (Nash & Newberg, 2023). Prior neurophysiological investigations of BKRY practitioners report frequency-specific changes, including decreased delta and increased theta–alpha activity (Sharma et al., 2018), increased theta indexing enhanced transitions from rest to meditation (Nair et al., 2017) and reorganisation of attention- and emotion-related networks (Sharma et al., 2023).

Building on this foundation, the present study aims to define the neurodynamic core of meditation through designing a methodologically robust experimental framework for systematic evaluation of two cardinal aims: 1) To identify meditation-specific EEG features that are distinct from other mental states, such as rest and a cognitive task, using broadband spectral power, aperiodic features and nonlinear EEG dynamics. 2) To assess the validity and reliability of the meditation-specific neuro-dynamic EEG features by studying their stability across four factors: a. Intra-subject/Inter-session; b. Inter-subject; c. Meditative states: guided vs. silent meditation; d. Meditation Expertise: Advanced vs. Intermediate.

## 2. Method

### 2.1 Participants

Thirty healthy Brahmakumaris Rajayoga (BKRY) meditators (20 males), aged 24-60 years with at least eight years (range: 7–27 years) or 3200 hours (range: 3200-32000 hours) of lifetime meditation experience, were recruited from various regional centres in Bangalore. For female participants, recordings were scheduled during the early follicular phase of their menstrual cycle (within 1 week of menstruation) to minimize the influence of hormonal fluctuations on brain activity (Avila-Varela et al., 2024). Before recruitment, permission for voluntary participation was obtained from the Spiritual Applications Research Centre (SpARC), Mt. Abu, India, thereby enabling meditators from different Brahmakumaris centres to participate in the study. Ethical approval was obtained from the institute’s Ethics Committee, where the study was conducted. All participants provided written informed consent before participation, and no incentives were offered.

The participants had a mean age of 41.1 ± 9.8 years. Of the 30 participants, nine were married; five were full-time householders; the remaining held private jobs, owned businesses, or were self-employed. Among the bachelors, three were full-time residents at different retreat centres, while the others worked in the following sectors: private (13), public (2), professional (2), and self-employed (1). Nearly 90% of participants had a strong educational background, either as graduates or having completed higher education, whereas only 3 had an education limited to the school level. Participants’ subjective meditation proficiency and self-reported progress in well-being were assessed using the BKRY Practice Proficiency Scale (Nair et al., 2018). The BKRY Practice Proficiency Scale measures an individual’s success in practising school-specific meditative thoughts. It consists of four subscales and 25 items, with a maximum score of 100. Only meditators who achieved a minimum score of 60, indicating a high level of subjective self-reported proficiency, were included in the study (Nair et al., 2018). On average, the participants reported 15.2 ± 6.3 years of experience practising BKRY meditation. Their mean score on the subjective practice proficiency scale was 81.5 ± 8.8.

### 2.2 Study Protocol

EEG signals were recorded for each subject under three conditions: meditation (M) with eyes open; rest (R) with eyes closed and open; and a working memory task (W). These conditions were administered in the following fixed sequence: R1-M1-R2—W—R3-M2-R4 (Figure 1).

**Fig. 1.**
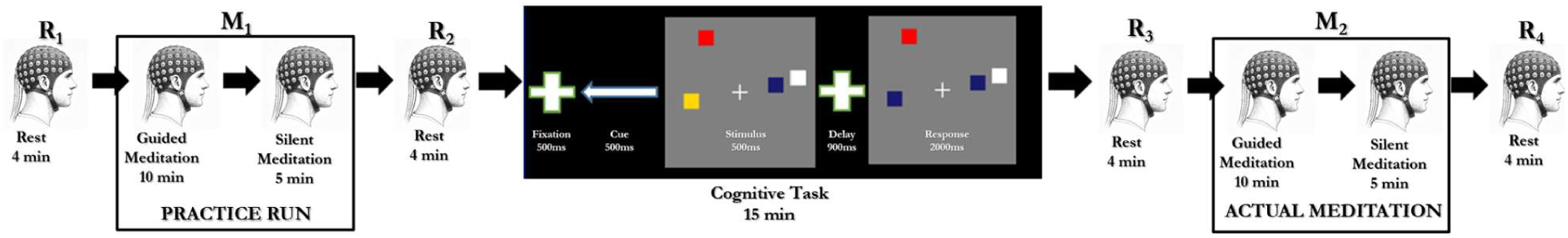
The arrow indicates the sequence of conditions in the protocol: R – rest (eyes open and closed, 2 minutes each); M – eyes-open meditation; W – cognitive task (working memory). The rest sessions correspond to R1 (pre-M1), R2 (post-M1), R3 (pre-M2), and R4 (post-M2) as described in the text.

The complete protocol, from R1 to R4, lasted approximately 70 minutes and was repeated on two consecutive days to examine the across-day reproducibility of the meditative state. To account for acclimatisation, the first meditation session (M1) on any day was treated as a practice run (Practice M), and the second session (M2) was considered the actual meditation session (Actual M). The working memory task (W) was positioned between the practice run and the actual meditation session to minimise any carryover effects from Practice M. Each meditation and working memory condition was preceded and followed by a rest condition.

The duration of each condition was as follows: Meditation (M): 15 minutes (10 minutes of guided meditation + 5 minutes of silent meditation). The guided meditation period consisted of a tradition-specific BKRY audio commentary in the participant’s preferred language (English, Hindi, or Kannada), delivered in the same voice and accompanied by soft background music. During the silent period, participants were asked to maintain their meditative state. Rest (R): 4 minutes, comprising 2 minutes with eyes closed and 2 minutes with eyes open. Participants were instructed to relax and refrain from meditating. Working Memory task (W): 15 minutes, participants performed a delayed matching task (Vogel & Machizawa, 2004) with arrays of coloured squares under mild-to-moderate cognitive load (set size: 2–3). They indicated, via button presses, whether the test stimuli matched the previously cued base stimuli. This task was chosen because it requires sustained, externally directed attention, in contrast to the inwardly focused attention characteristic of meditation. All stimuli and instructions were delivered using E-Prime 2.0 (Psychology Software Tools, Inc., Sharpsburg, PA, USA) on a 34 × 27 cm LCD monitor placed 90 cm from the participant.

### 2.3 Data acquisition

EEG recordings were obtained using a 64-channel ActiCHamp Plus amplifier with Brain Recorder 1.23 software (Brain Products GmbH, Germany) and an actiCAP active electrode system following the 10/10 international placement system. Signals were sampled at 10 kHz with default hardware filters (DC to Nyquist frequency), and FCz was used as the online reference electrode. Electrode impedances were maintained below 50 kΩ, in accordance with the manufacturer’s recommendations for active electrode systems. Recordings were consistently scheduled between 9:00 a.m. and 1:00 p.m. for all participants. For each participant, sessions on both days were conducted at approximately the same time to minimise circadian effects on EEG signals. Participants were seated in a comfortable chair in a dimly lit, sound-attenuated recording room, with the temperature maintained at 25 °C and the relative humidity between 40% and 60%.

## 3. Data analysis

### 3.1 Pre-processing of EEG data

EEG data were preprocessed using EEGLAB v2024.0 (Delorme & Makeig, 2004) and custom MATLAB scripts (R2023b). The pre-processing steps were as follows:

1. Resampling and Filtering Continuous EEG data were down-sampled to 250 Hz, band-pass filtered between 0.5 and 120 Hz, and a band-stop filter was applied at 48–52 Hz to remove line noise.
2. Channel Cleaning Noisy channels were identified using the *clean_rawdata* plugin and replaced via spherical spline interpolation.
3. Independent Component Analysis (ICA), Artifact Removal & re-referencing An initial ICA (Infomax algorithm) (Bell & Sejnowski, 1995) was conducted on a temporary dataset containing eye blinks (2-second epochs around blinks created using Brainstorm functions). This was necessary because the participants, habituated to open-eye meditation, exhibited fewer, more spaced blinks, reducing the effectiveness of automatic ICA blink detection. Blink-related components identified using ICALabel were removed, and the cleaned data were then recombined. On this blink-corrected continuous data, Artifact Subspace Reconstruction (ASR) with a threshold of 20 standard deviations was used to remove transient bad segments. A second ICA was performed on the ASR-corrected data to identify and remove other artifact components (e.g., muscle, cardiac, residual line noise), again using ICALabel. After ICA-based correction, a final ASR step was applied at the 2-s epoch level to ensure removal of any residual artifacts. Data were then re-referenced to a common average reference.
4. Epoching and Feature Extraction Cleaned continuous EEG data were segmented into 4-second epochs with 50% overlap for feature extraction. Table 1 provides a detailed description of the EEG measures included in the study Broadband Spectral features: Relative power was computed using Welch’s method for the following bands: delta (1–4 Hz), theta (4–8 Hz), theta–alpha (6–10 Hz), alpha (8–12 Hz), lower beta (12–18 Hz), higher beta (18–30 Hz), and gamma (30–80 Hz). The additional theta–alpha (Sharma et al., 2023) and higher beta bands were included based on prior meditation studies (Cahn & Polich, 2006; Thomas et al., 2014). Aperiodic features: The slope and intercept of the 1/f component were extracted using FOOOF and IRASA methods (Donoghue et al., 2020; Gerster et al., 2022) with their popular implementations in two Python libraries (https://fooof-tools.github.io/fooof/; https://yasa-sleep.org/generated/yasa.irasa.html). Nonlinear features: These were estimated using the Python *Antropy* library, including Higuchi’s fractal dimension (Higuchi), Detrended Fluctuation Analysis (DFA) (Uudeberg et al., 2025) and Permutation Entropy (Y. Han et al., 2020).

**Table 1:** Detailed description of EEG measures assessed in this study.

| EEG-based Measures | Feature | Quantifies | Functional Interpretation | Prior findings from meditation studies |
| --- | --- | --- | --- | --- |
| Broadband Spectral Features | Delta (1-4 Hz) | Spectral redistribution of power within canonical frequency bands (Wen & Liu, 2016) | Characterises deep sleep, cortical inhibition; In waking states reflect attentional engagement | ↓ Delta reported in advanced meditators during deep absorption states with no drowsiness; ↑ during meditation, if asked to focus on an external engaging task (Cahn et al., 2010, 2013) |
|  | Theta (4-8 Hz) |  | Internal attention, memory, cognitive control, interoceptive awareness | ↑ frontal-midline theta (Aftanas & Golocheikine, 2001; Braboszcz et al., 2017; Cahn et al., 2013) during focused attention and mindfulness meditation (Lomas et al., 2015). Often stronger in experienced meditators; can reflect both effortful control (novices) and stable attentional engagement (experts) (Cahn & Polich, 2006). |
|  | Theta-Alpha (6-10 Hz) |  | Balance between internal (theta) and relaxed alertness (alpha); attentional stability | State-level ↑ in theta–alpha, especially during sustained attention practices (Panda et al., 2016; Sharma et al., 2023). Less studied |
|  | Alpha (8-12 Hz) |  | Relaxed alertness, sensory inhibition, top–down regulation | Robust ↑ alpha power and coherence during meditation, especially in posterior regions, reflect calm awareness (Braboszcz et al., 2017; Cahn et al., 2013; Fell et al., 2010; Lee et al., 2018; Travis & Shear, 2010) |
|  | Low-Beta (12-18 Hz) |  | Active cognition, alertness, sustained attention | Mixed findings: with reports of no change, increases, or decreases depending on the type of meditation (Faber et al., 2015; Lee et al., 2018) |
|  | High-Beta (18-30 Hz) |  | Arousal, motor readiness, anxiety, cognitive effort | Often ↓ during meditation, particularly in experienced practitioners (DeLosAngeles et al., 2016; Thomas et al., 2014) |
|  | Gamma (30-80 Hz) |  | Sensory and cognitive processes, integration, conscious awareness | ↑ Gamma power and synchrony reported in long-term practitioners (Braboszcz et al., 2017; Cahn et al., 2010; Lutz et al., 2004) |
| Aperiodic Features | Aperiodic Intercept (1/f offset) | Overall broadband power of the 1/f background noise in log-power space. | Global neural activity level; population firing intensity | Scarcely studied |
| | Aperiodic Slope (1/f exponent) | Steepness of power decay across frequencies | Balance between excitation and inhibition (E/I balance); neural noise | Flatter slopes (↓ exponent) reported during focused and advanced meditation, suggesting increased cortical excitability (McQueen et al., 2024; Pascarella et al., 2025) |
| Nonlinear measures | DFA (Hardstone et al., 2012) | Long-range temporal correlations in non-stationary physiological time series | Temporal integration; self-organised criticality | Band-wise reduction of DFA (Irmischer et al., 2018) ↓DFA during meditation suggests enhanced ability to handle complexity (Hirekhan et al., 2019) ↑DFA reported in calming meditation (Vyšata et al., 2014) |
| | Higuchi (Higuchi, 1988) | Signal complexity in the time domain through measurement of the waveform's length at multiple scales and analysing how this length changes with scale | Irregularity and dimensionality of neural dynamics | ↑ Higuchi during meditation (Walter & Hinterberger, 2022a; Young et al., 2021) ↑Higuchi in novice and advanced meditators (Teachers), no change in advanced meditators (Senior practitioners) (Kakumanu et al., 2018) |
| | Permutation Entropy (PE) (Bandt, 2002) | Local temporal unpredictability of the signal | Information complexity and randomness | ↑ PE during meditation compared to rest or task states (Atad et al., 2025; Walter & Hinterberger, 2022b) ↓PE reported in calming and insight meditation (Vyšata et al., 2014) |

### 3.2 Statistical comparison

#### 3.2.1 Distinguishing meditation from other mental states

##### 3.2.1.1 At the electrode level - Topographical comparisons

To analyse scalp distribution of differences between three mental states, electrode-wise LIMO EEG analyses were conducted using the LIMO toolbox (Pernet et al., 2011). LIMO performs robust t-statistics with 2000 permutations and adjusts for multiple comparisons across electrodes using the threshold-free cluster enhancement (tfce) method (Pernet et al., 2015). We chose tfce over traditional clustering because it increases t-values for each electrode by aggregating cluster mass across all thresholds, resulting in electrode-specific tfce scores. To control the family-wise error rate (FWER), the maximum tfce scores were permuted to obtain corrected p-values. The significance threshold was set at p < 0.05, and the maximum t-value for a significant effect was reported.

##### 3.2.1.2 Global Electrode Average -

To determine meditation-specific EEG measures distinct from other mental states, EEG features were compared between meditation and rest, and between meditation and task, using robust permutation-based repeated-measures (RM) ANOVA. Post hoc comparisons were performed using paired Wilcoxon signed-rank tests with Bonferroni corrections.

#### 3.2.2 Validity and reliability assessment of meditation-specific features

##### 3.2.2.1 Intra-subject reliability

Intra-subject or inter-session reliability across each feature was assessed using the Intra-class Correlation Coefficient (ICC), a widely used reliability metric that quantifies the consistency of repeated measurements (Bruton et al., 2000). ICC estimates, and their 95% confidence intervals (CI) were calculated using RStudio scripts (Posit Team, 2025) based on mean measurements, absolute agreement, and a two-way mixed-effects model—specifically ICC(2,k) (McGraw & Wong, 1996; Shrout & Fleiss, 1979). Interpretation was based on the CI of ICC estimates (Koo & Li, 2016): a) Excellent: lower and upper limits > 0.90; b) Good: both limits between 0.75 and 0.90; c) Moderate: both limits between 0.50 and 0.75; d) Poor: both limits < 0.50. Repeatability was assessed by comparing measurements taken on Day 1 and Day 2.

##### 3.2.2.2 Inter-subject reliability

To examine inter-subject stability of meditation-specific features, inter-subject correlation (ISC) and inter-subject distance (ISD) were computed for each day by creating a multivariate feature space of meditation-specific features. Analyses were done separately for the silent and guided meditation periods of the Actual M session on each recording day. ISC and ISD were computed using Pearson’s correlation coefficient and Euclidean distance, respectively. To this end, for each subject, a vector of multivariate features was created, and each subject’s vector was channel-wise compared with another subject’s vector, yielding a symmetric N x N matrix for inter-subject correlation and inter-subject Euclidean distance.

##### 3.2.2.3 Validity across meditative states and meditation experience

Stability or variability across meditative states (guided versus silent meditation) and across expertise (advanced versus intermediate meditators) was assessed using Yuen’s robust t-test with 20% trimmed means across features.

## 4 Results

The results are organised into two sections. The first section presents meditation-specific EEG features that are distinct from those observed under baseline resting and task conditions. This is achieved by comparing the meditative state with other mental states at two levels: topographical and global average. The second section examines the validity and reliability of the meditation-specific features by assessing their stability across four factors: over time (intra–subject stability), across subjects (inter–subject stability), across meditative states (guided vs. silent meditation), and across experience (meditation expertise).

### 4.1. Meditation-specific neuronal (EEG) measure

#### 4.1.1. Topographical Comparison

For analysis at the topographical level, electrode-wise EEG feature values averaged across all meditation sessions over two days, comprising both guided and silent states, were compared with those of baseline eyes-open rest and task conditions, using robust t-statistics in the LIMO toolbox (Supplementary file: Figures 1 and 2).

##### 4.1.1.1. Broadband spectral features

On comparing conditions for broadband spectral power for each frequency band, meditation showed significantly reduced relative power in the delta band (frontoparietal electrodes; max t <-4.9, p < 0.05) and the low-beta band (centroparietal electrodes; max t <-5.9, p < 0.05). compared to rest. Compared with the task condition, meditation showed reduced relative power in delta (widespread electrodes; max t ≤ - 8.02, p < 0.01), low beta (frontoparietal electrodes; max t ≤ -4.7, p < 0.05), high beta (parietal electrodes; max t ≤-3.3, p < 0.05), and gamma bands (centroparietal electrodes; max t ≤-4.8, p < 0.05). Conversely, meditation showed significantly increased power in theta (centroparietal electrodes; max t ≤5.2, p < 0.05) and theta-alpha bands (centroparietal and few peripheral electrodes; max t ≤ 4.47, p < 0.05) relative to rest, and elevated power in theta-alpha (widespread electrodes; max t ≤ 5.24, p < 0.01) and alpha bands (widespread electrodes; max t ≤6.24, p < 0.001) compared to task.

##### 4.1.1.2. Aperiodic features

In aperiodic features, Intercept IRASA (widespread electrodes; max t ≤ 5.56, p < 0.05) was found to be increased, while Slope FOOOF (frontocentral electrodes; max t ≤ -4, p < 0.05) showed decreases during meditation compared to the task. A similar trend was observed in comparisons between meditation and rest, with increased Intercept IRASA (widespread electrodes; max t ≤ 14.05, p < 0.0001) and reduced Slope FOOOF (frontocentral electrodes; max t ≤ -4.5, p < 0.05) during meditation. In addition, Slope IRASA (widespread electrodes; max t < -14.6; p < 0.0001) showed a significant decrease during meditation compared with rest.

##### 4.1.1.3. Nonlinear features

DFA was found to be increased during meditation compared to both rest (widespread electrodes; max t ≤ 5.3, p < 0.05) and task (widespread electrodes; max t ≤ 5.63, p < 0.05) conditions. Whereas, Higuchi and Permutation entropy were found to reduce during meditation in comparison to both rest (widespread electrodes; max t ≤ -8.5, p < 0.001; widespread electrodes; max t ≤ -33.7, p < 0.0001) and task (widespread electrodes; max t ≤ -6.9, p < 0.05; widespread electrodes; max t ≤-6.2, p < 0.05) conditions respectively.

#### 4.1.2. Global Average Comparison

Robust Permutation RM ANOVA was used to compare the global average values across three conditions for each EEG feature. Post hoc comparisons were performed using paired Wilcoxon signed-rank tests, with p-values adjusted for multiple comparisons using the Bonferroni correction (Figure 2.1). Following EEG features showed significant condition effects (Table 1) with Permutation Entropy amongst nonlinear features showing highest significant effect size (F = 54.28, p < .001, η²p = 0.80), followed by Intercept IRASA (F = 37.02, p < .001, η²p = 0.67) and Slope IRASA among aperiodic features (F = 27.81, p < .001, η²p = 0.57), Higuchi among nonlinear features (F = 20.51, p < .001, η²p = 0.34), alpha (F = 35.56, p < .001, η²p = 0.40), theta–alpha (F = 25.77, p < .001, η²p = 0.28), delta (F = 26.08, p < .001, η²p = 0.27), and high beta bands (F = 17.8, p < .001, η²p = 0.22) amongst spectral features and DFA (F = 10.01, p < .001, η²p = 0.22) being the least significant from nonlinear features. Post hoc paired Wilcox comparisons (Bonferroni corrected) revealed a significant increase in theta-alpha power and DFA and a significant reduction in Permutation Entropy and Higuchi during meditation compared with both rest (p < .05) and task (p < .01). Table 2 presents the F-statistic, effect size, and post hoc comparisons for each EEG feature.

**Fig. 2.1.**
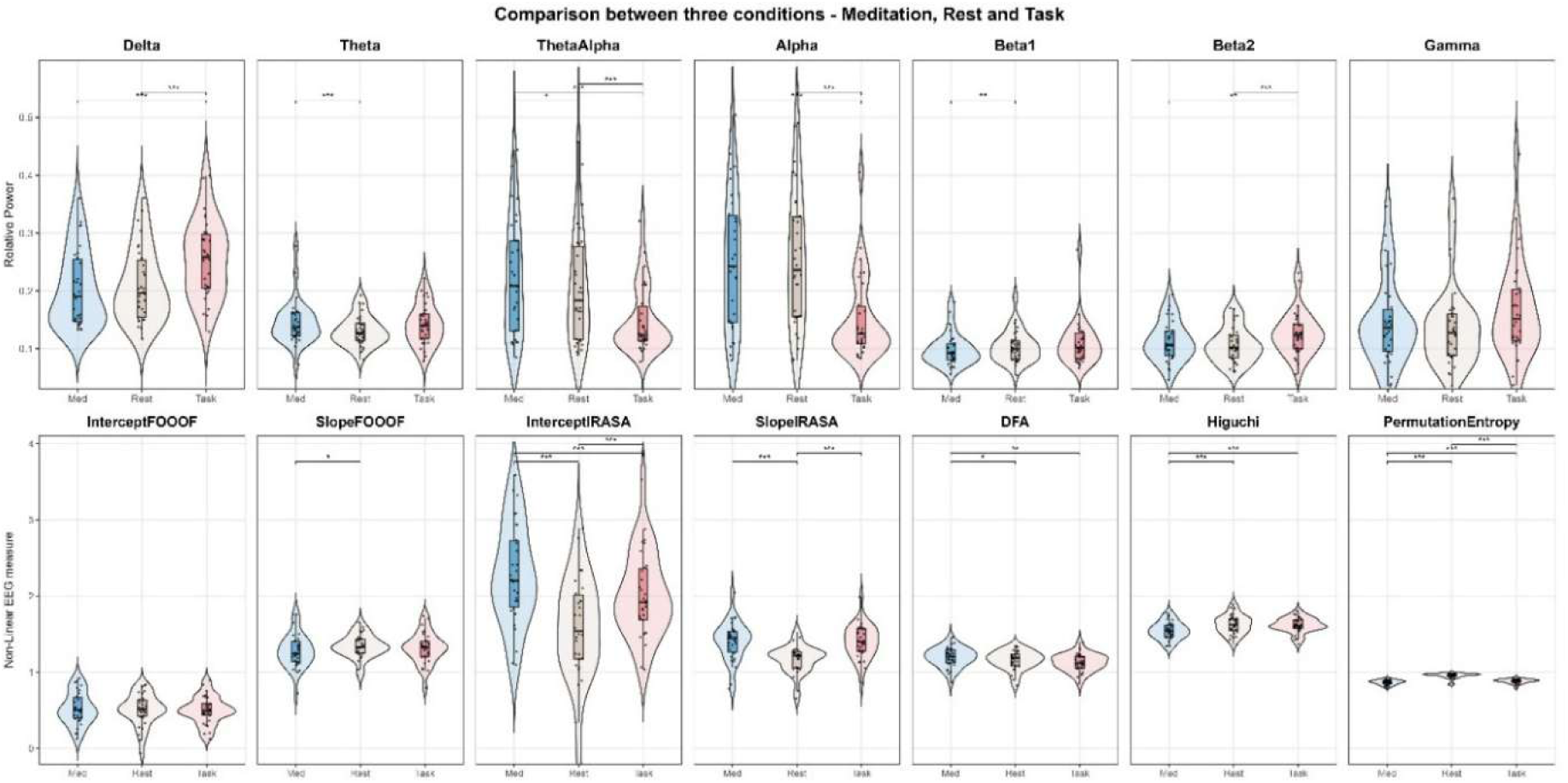
Comparison between Meditation, Rest and Task using Robust Permutation RM ANOVA and post-hoc comparisons using paired Wilcoxon signed-rank tests and adjusted p (Bonferroni corrected). Med – all Meditation sessions, including guided and silent meditation, averaged; Rest – Average of eyes open baseline rest on two days; Task – Average of the working memory task on two days. The top panel compares three conditions with respect to spectral features (Relative power), while the bottom panel depicts Aperiodic and Nonlinear EEG Features.

**Table 2:** Comparison between Meditation, Rest and Task using Robust Permutation RM ANOVA and post-hoc comparisons using paired Wilcoxon signed-rank tests and adjusted p (Bonferroni corrected)

| EG Features | | F Statistic | $\eta^2 p_{\text{partial}}$ | Post-hoc comparisons | | |
| --- | --- | --- | --- | --- | --- | --- |
|  |  |  |  | M - R | M - W | R - W |
| Broadband Spectral Features | Delta | 26.08*** | 0.27 | NS | *** | *** |
|  | Theta | 7.45* | 0.12 | *** | NS | NS |
|  | ThetaAlpha | 25.77*** | 0.28 | * | *** | *** |
|  | Alpha | 35.56*** | 0.40 | NS | *** | *** |
|  | LowBeta | 5.7* | 0.10 | ** | NS | NS |
|  | HighBeta | 17.8** | 0.22 | NS | ** | *** |
|  | Gamma | 4.9 | 0.12 | NS | NS | NS |
| Aperiodic Features | Intercept FOOOF | 0.91 | 0.01 | NS | NS | NS |
|  | Slope FOOOF | 4.3 | 0.09 | * | NS | NS |
|  | Intercept IRASA | 37.02*** | 0.67 | *** | *** | *** |
|  | Slope IRASA | 27.81*** | 0.57 | *** | NS | *** |
| Non-linear Features | DFA | 10.01** | 0.22 | * | ** | NS |
|  | Higuchi | 20.51*** | 0.34 | *** | *** | NS |
|  | Permutation Entropy | 54.28*** | 0.80 | *** | *** | *** |

M – all Meditation sessions, including guided and silent meditation, averaged; R – Average of eyes open baseline rest on two days; W – Average of the working memory task on two days. Statistical significance (p) is denoted by \* $p < 0.05$ , \*\* $p < 0.01$ , \*\*\* $p < 0.001$ . In the post hoc comparison, a significant difference is indicated by \*, NS – Nothing significant.

##### Meditation-specific EEG features distinct from both rest and task in Topographical and Global Average Comparisons

Theta-alpha band amongst spectral features, Intercept IRASA amongst aperiodic and all nonlinear measures (DFA, Higuchi and Permutation Entropy) were distinct in meditation from both rest and task conditions in both comparisons (Figure 2.2).

**Fig. 2.2.**
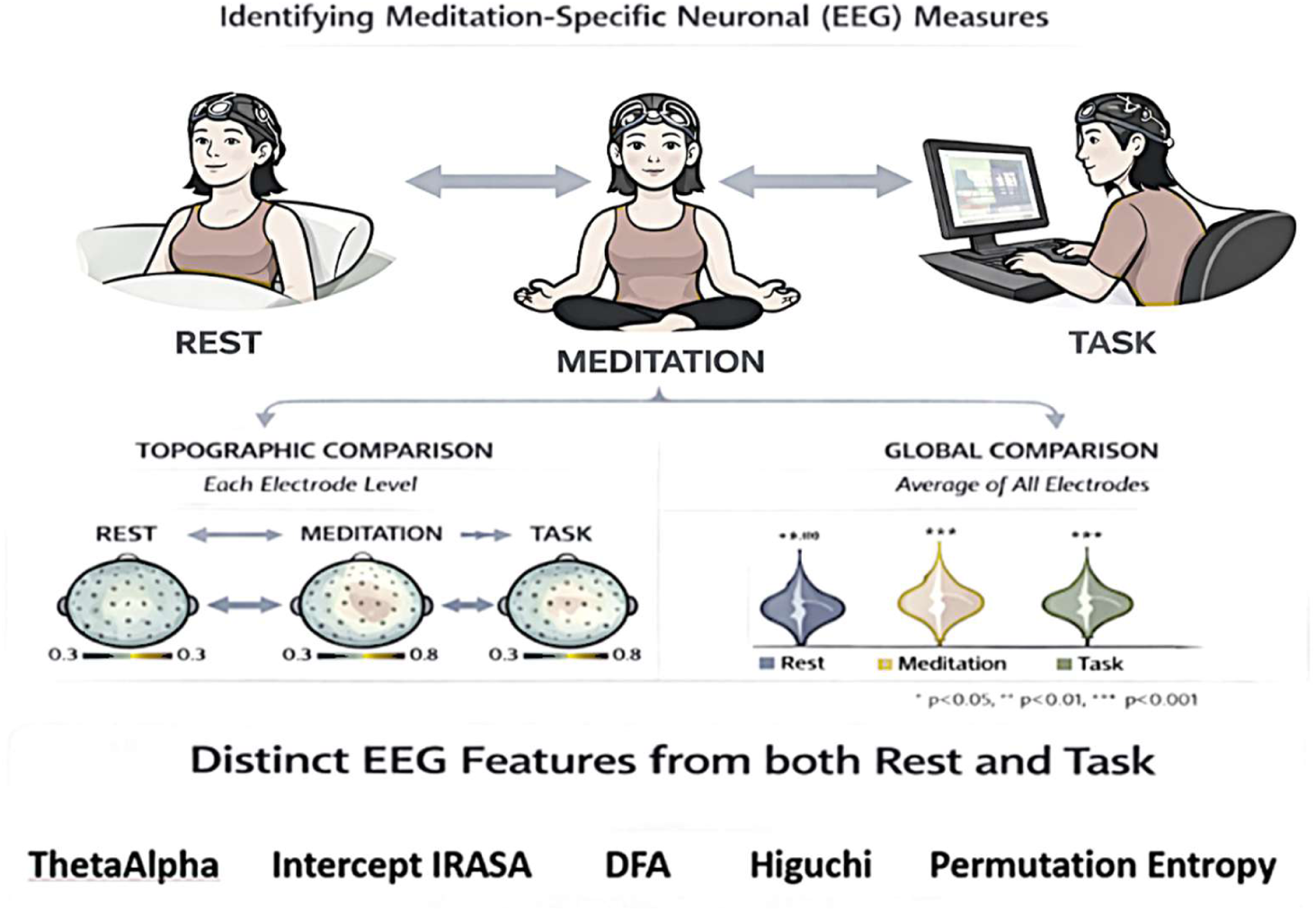
Meditation-specific Neuronal (EEG) measures distinct from both rest and task conditions

### 4.2. Validity and reliability of the meditation-specific EEG features

#### 4.2.1. Intra-subject stability

ICC was used to assess within-subject stability of meditation-specific EEG features over time (across days) during the actual meditation session for both silent and guided meditation periods (Figure 3). Both silent and guided meditation demonstrated good to excellent reliability (ICC > 0.89), with theta-alpha showing the greatest stability (CI limits > 0.9). Guided meditation showed excellent reliability (ICC > 0.92) across all features, except for Permutation Entropy (Table 3).

**Fig. 3.**
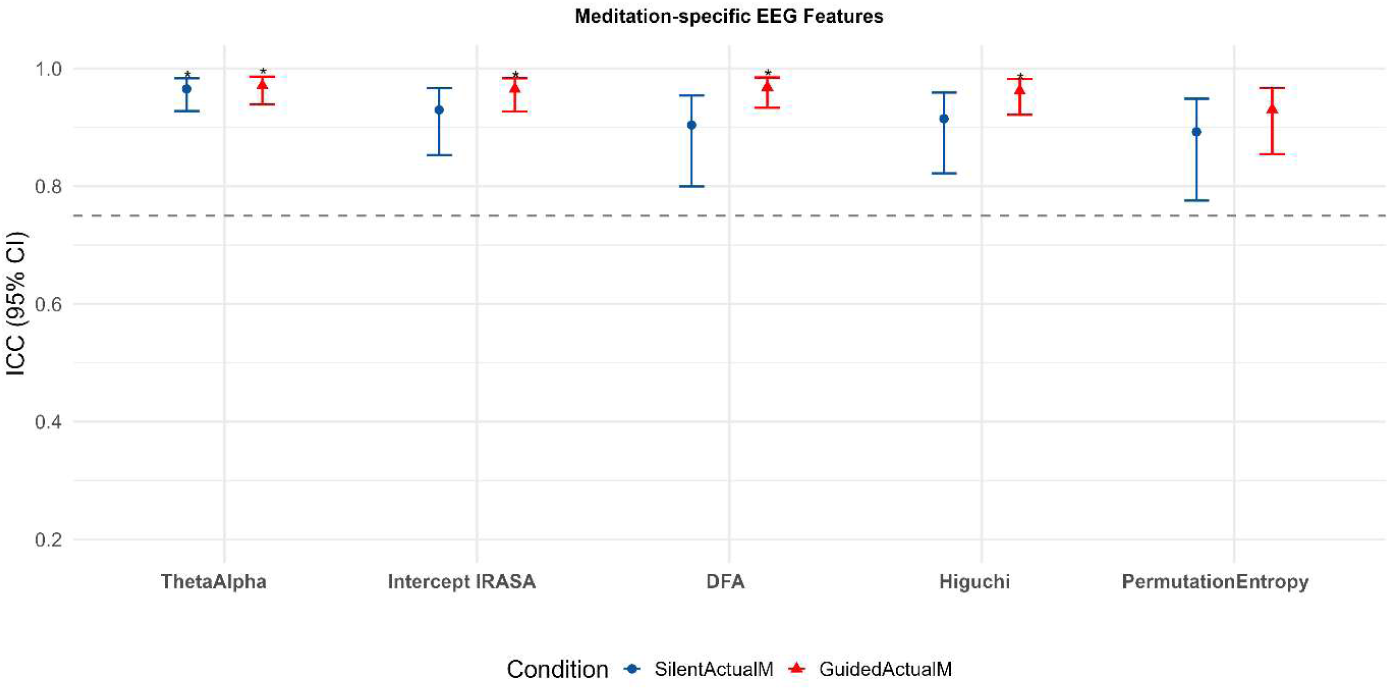
Intra-subject stability of meditation-specific EEG features during the silent and guided meditation periods of Actual M session across days

**Table 3:** ICC estimates with CI limits for each of the Meditation-specific EEG features during the silent and guided meditation periods of the Actual M session across days.

| EEG Features | EEG Variable | ICC Value [CI - Lower limit, CI - Upper limit] |  |
| --- | --- | --- | --- |
|  |  | For Actual Meditation sessions pairs |  |
|  |  | Between Silent Actual<br>M Sessions | Between Guided<br>Actual M Sessions |
| Broadband Spectral<br>Feature | Theta-Alpha | 0.97** [0.93,0.98] | 0.97** [0.94, 0.99] |
| Aperiodic Feature | IRASA Intercept | 0.93* [0.85,0.97] | 0.97** [0.93, 0.98] |
| Nonlinear Features | DFA | 0.90* [0.80,0.95] | 0.97** [0.93, 0.98] |
|  | Higuchi Complexity | 0.91* [0.82,0.96] | 0.96** [0.92, 0.98] |
|  | Permutation Entropy | 0.89* [0.78,0.95] | 0.93* [0.85, 0.97] |

\* Signifies good to excellent repeatability; \*\* signifies excellent repeatability

#### 4.2.2. Inter-subject stability

To assess Inter-subject reliability, Inter-subject correlation (ISC) and Inter-subject distance (ISD) were computed for the multivariate feature (meditation-specific) between all pairs of subjects, yielding separate matrices for ISC and ISD. Pairwise ISC matrices revealed high correlations between subjects for both silent and guided meditation on days 1 and 2 (Figures 4.1 & 4.2, top-left). Pairwise ISD heatmaps (Figures 4.1 & 4.2, top-right) showed moderate inter-subject dispersion across the two meditation periods on both days, with no strong clustering patterns. For further statistical analysis, paired Wilcoxon signed-rank tests were performed on subject-wise mean ISC/ISD values across days, with false discovery rate correction applied across domains. A significant increase in mean ISC from day 1 to day 2 (p < 0.001) and a significant reduction in mean ISD on day 2 relative to day 1 (p < 0.001) were observed only in silent meditation (Figure 4.1, bottom panel).

**Fig. 4.1.**
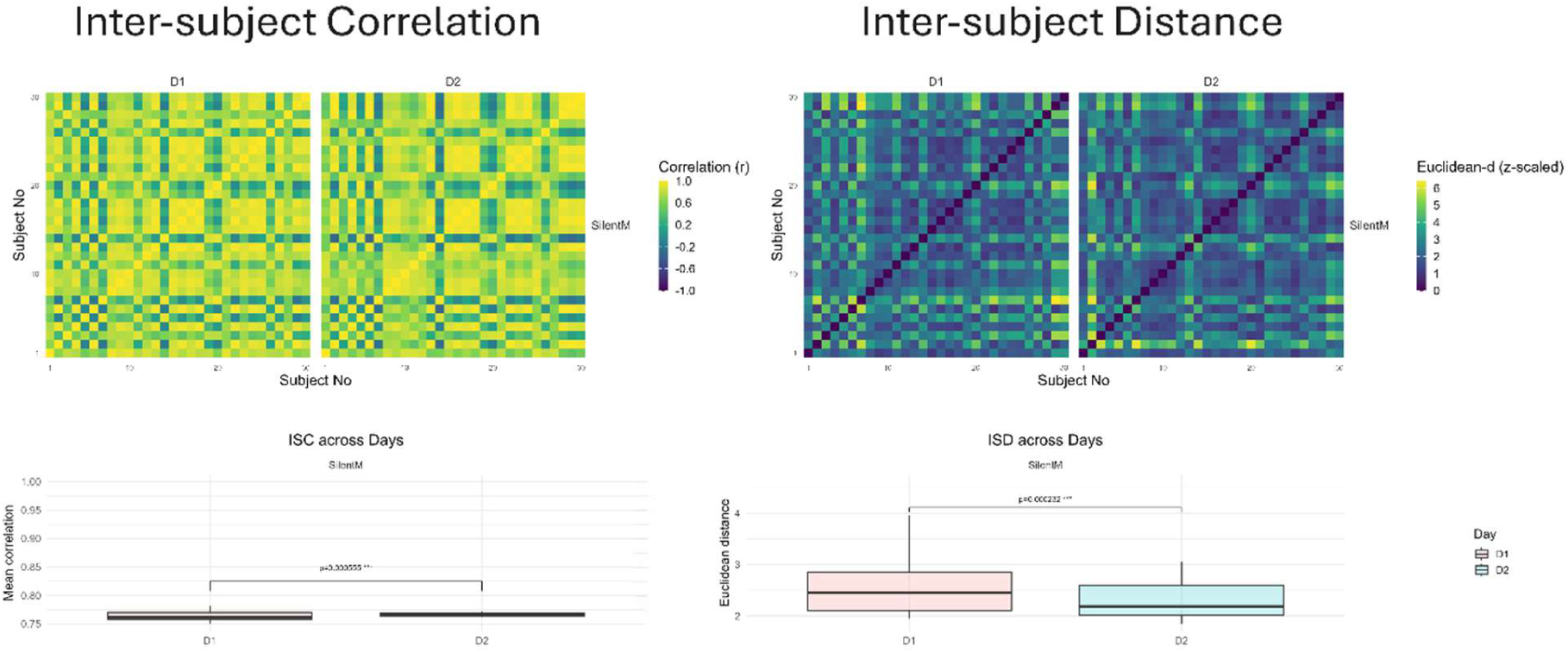
Inter-subject stability of meditation-specific EEG features during the silent meditation period of the Actual M session across days Top Row: Inter-subject correlation (ISC, left) and Inter-subject distance (ISD, right) matrices computed from multivariate EEG features for day 1 (D1) and day 2 (D2). Each cell represents pairwise similarity or distance between subjects. Bottom row: Subject-wise means ISC and ISD across days. Boxplots show median and interquartile range; brackets indicate paired comparisons across days with FDR-corrected *p*-values.

**Fig. 4.2.**
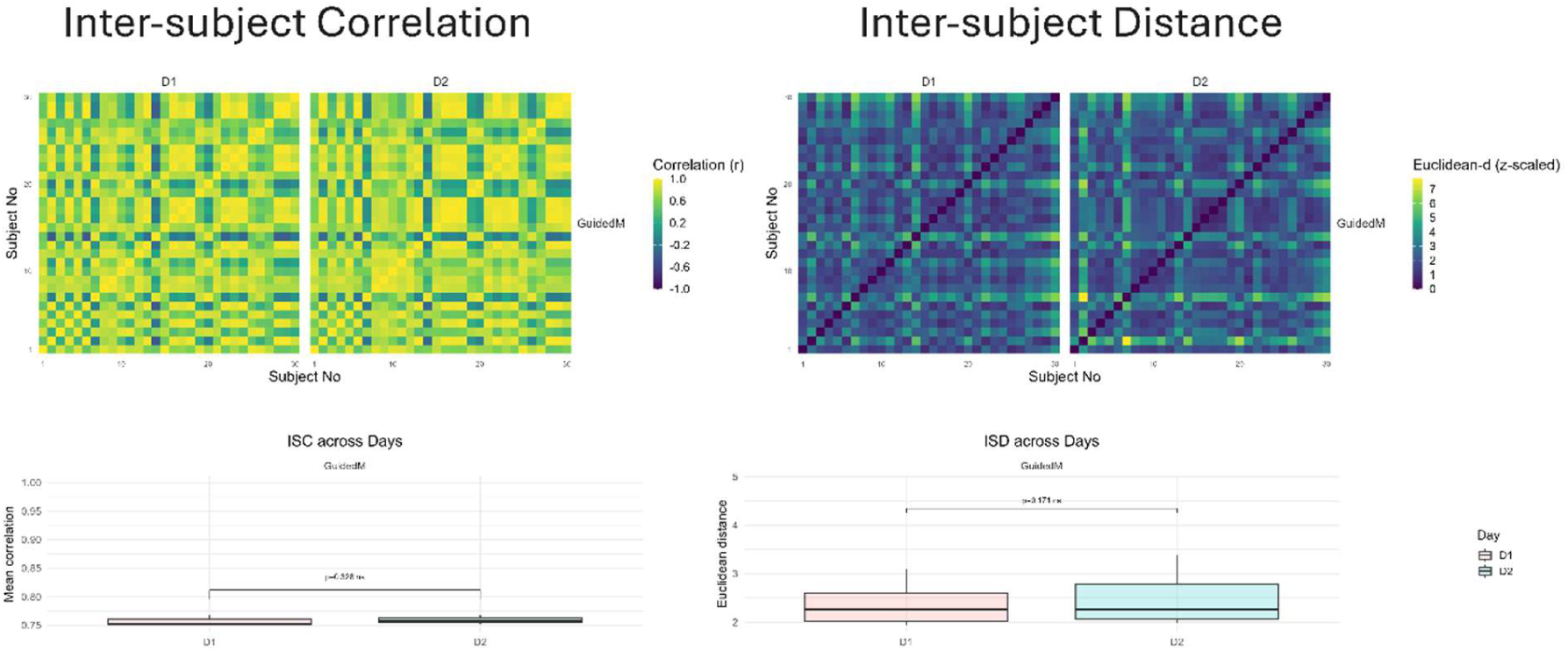
Inter-subject stability of meditation-specific EEG features during the guided meditation period of the Actual M session across days Top Row: Inter-subject correlation (ISC, left) and Inter-subject distance (ISD, right) matrices computed from multivariate EEG features for day 1 (D1) and day 2 (D2). Each cell represents pairwise similarity or distance between subjects. Bottom row: Subject-wise means ISC and ISD across days. Boxplots show median and interquartile range; brackets indicate paired comparisons across days with FDR-corrected *p*-values.

#### 4.2.3. Guided vs Silent meditation

To assess the differences for meditation-specific features across guided and silent meditation periods of the Actual M session, Yuen’s robust t-test (20% trimmed means) with FDR correction across features was performed. Visualisation using raincloud plots revealed that none of the EEG features differs significantly between guided and silent meditation on either day (Figure 5).

**Fig. 5.**
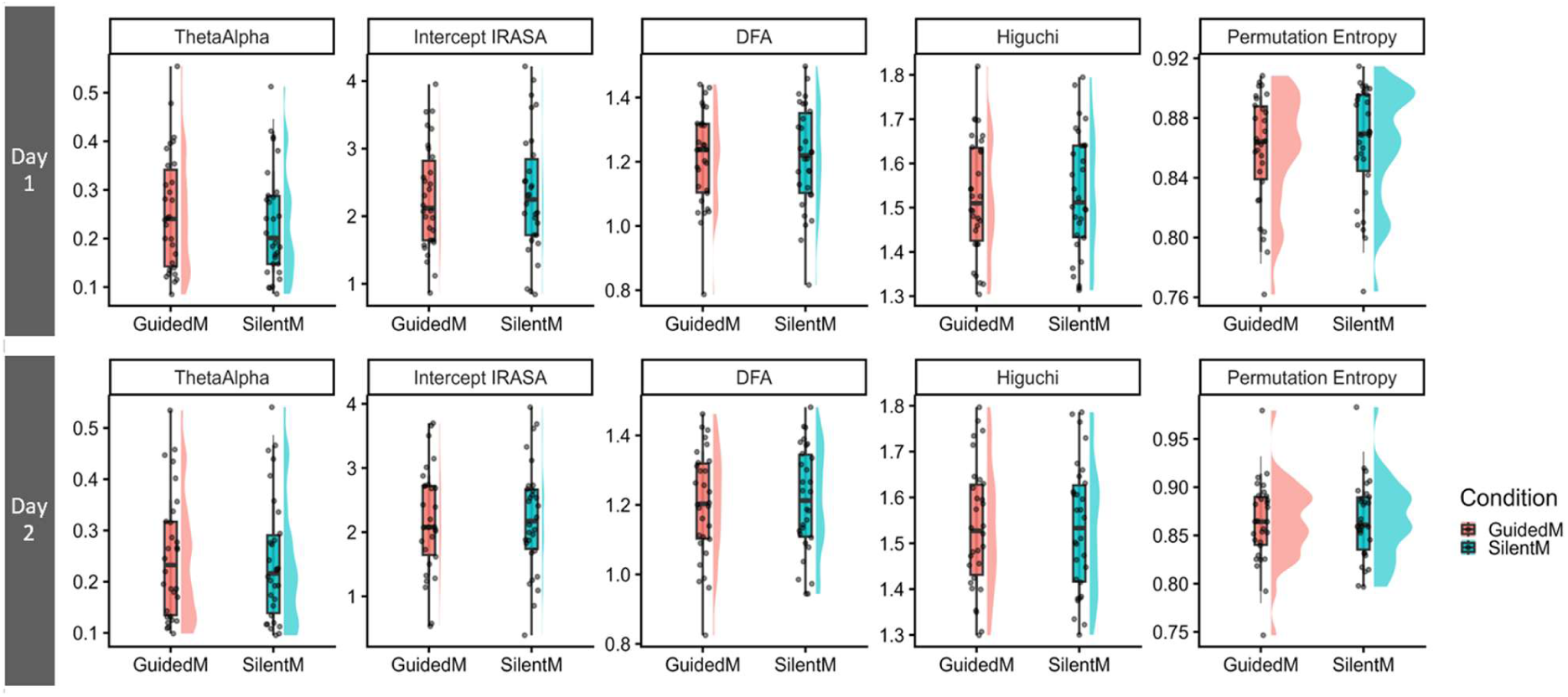
Raincloud plots with Yuen’s robust t-test (20% trimmed means) with FDR correction to assess differences between Guided and Silent periods of the Actual M session

#### 4.2.4. Meditation expertise-based differences

To assess if the meditation-specific features differed with meditation expertise based on the number of hours of practice. All features were compared between Advanced (> 10000 hours) and Intermediate meditators (< 10000 hours) on day 1 and day 2 for both the guided and silent periods of the Actual M session, using Yuen’s robust t-test (20% trimmed means) with FDR correction. Visualisation using box plots revealed no significant differences in any EEG features between the two meditator groups on either day, during both guided and silent meditation periods (Figure 6).

**Fig. 6.**
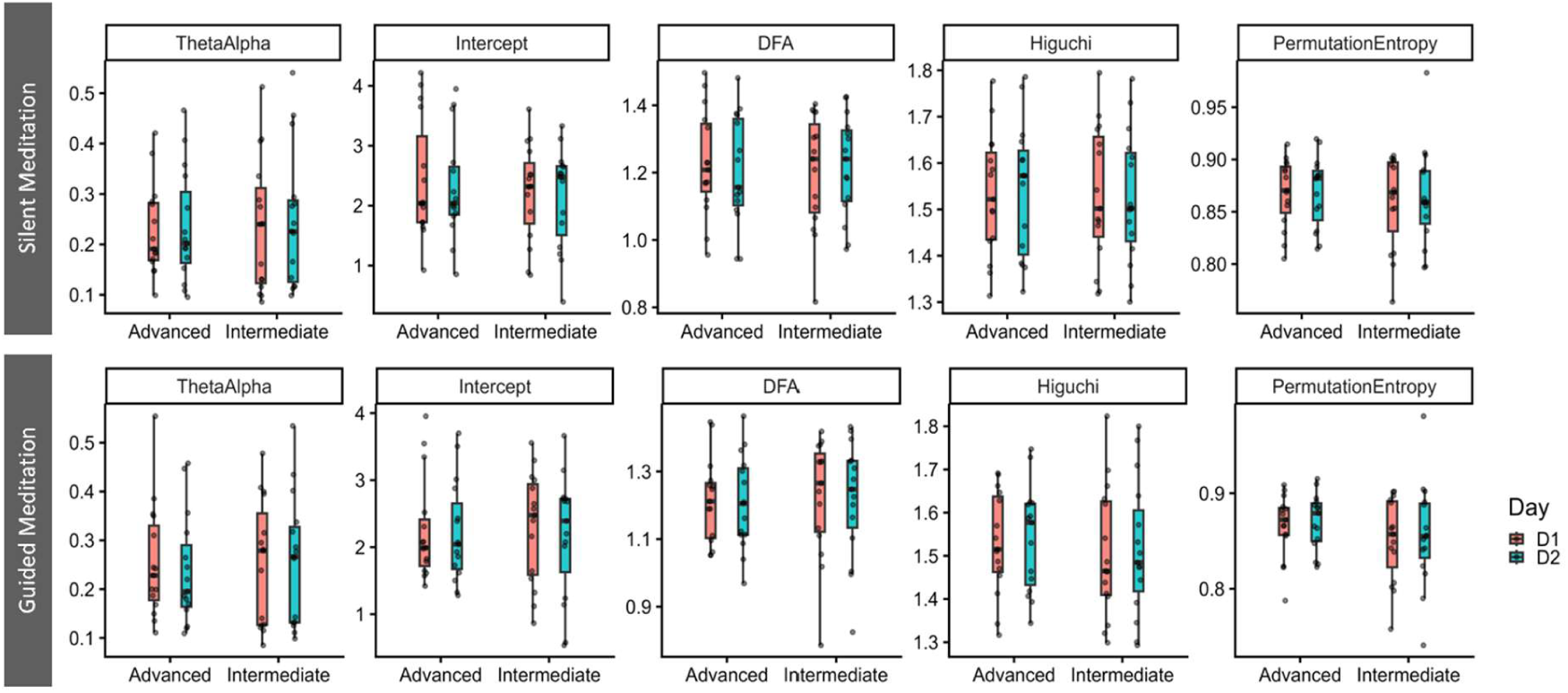
Raincloud plots with Yuen’s robust t-test (20% trimmed means) with FDR correction to assess differences between Advanced and Intermediate meditators, during guided and silent periods of Actual M session

## 5 Discussion

This study implemented a reliability-informed experimental protocol to systematically examine broadband spectral, aperiodic, and nonlinear EEG features in experienced long-term meditators. By contrasting meditation with both passive rest and an active cognitive task, we identified a meditation-specific neurodynamic core distinct from adjacent mental states. Importantly, we evaluated the validity and reliability of these features across four critical dimensions: intra-subject (inter-session) stability, inter-subject consistency, meditative format (guided vs. silent), and practitioner expertise (advanced vs. intermediate). Our findings demonstrate that meditation-specific neural markers constitute stable and reproducible signatures in experienced practitioners, independent of session repetition, inter-individual variability, meditation format, or expertise level (Figure 7).

**Fig. 7.**
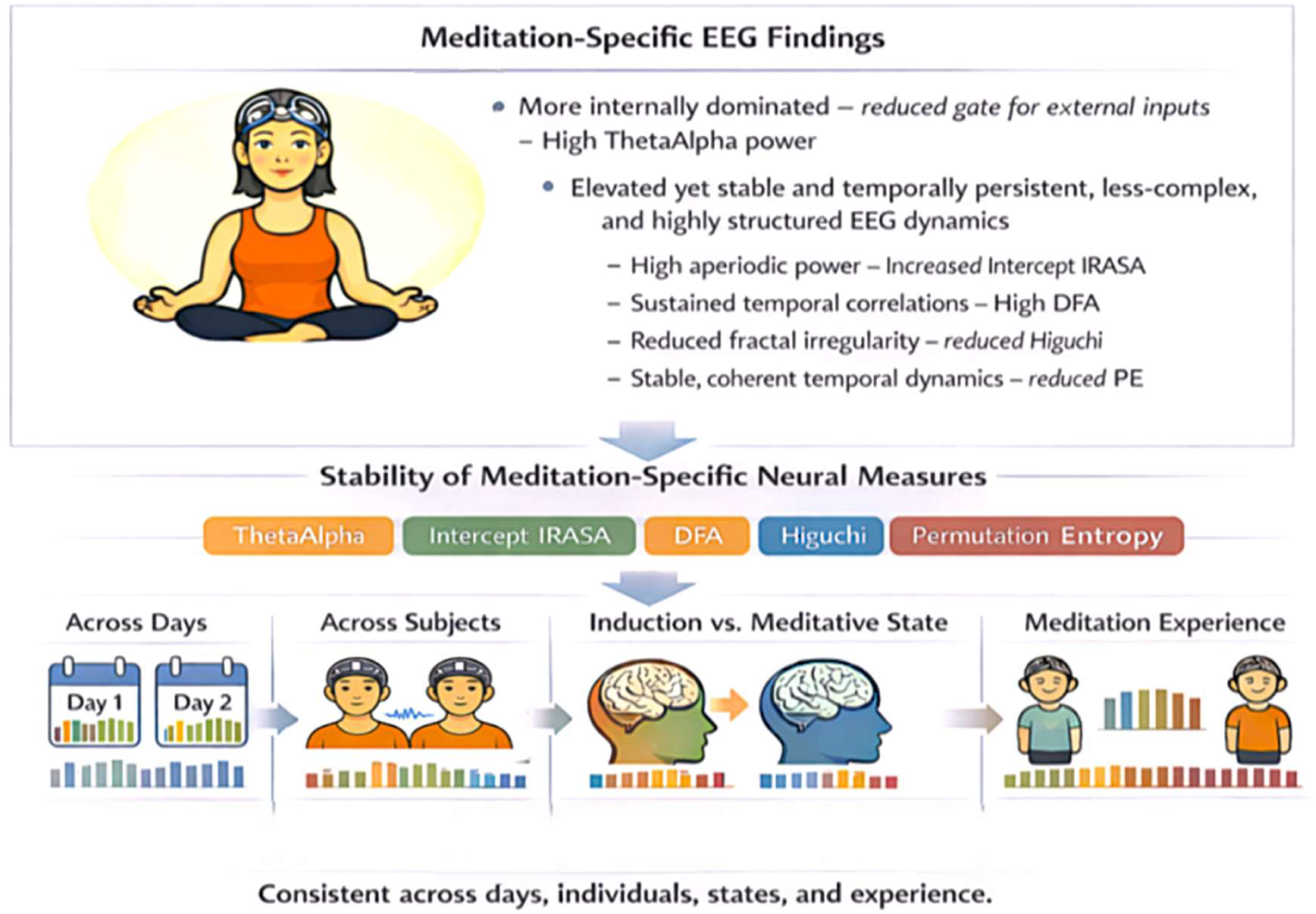
Findings summary of the study

### 5.1 Meditation-specific neural correlates distinct from other mental states

Most EEG studies infer meditation-specific neural correlates by comparing experienced meditators with novices (Baijal & Srinivasan, 2010; Braboszcz et al., 2017; Katyal & Goldin, 2021) or between practitioners of varying expertise (Kakumanu et al., 2018; Nair et al., 2017), or through comparisons with the resting baseline (Cahn et al., 2010; Sharma et al., 2018, 2023). Few incorporate alternative cognitive states as active comparators (DeLosAngeles et al., 2016; Malipeddi et al., 2024; Thomas et al., 2014). By including both rest and task conditions, we tested whether meditation exhibits distinct neurodynamic properties beyond passive baseline effects and active cognitive engagement. Across topographical and global-average analyses, five features emerged as core markers distinguishing meditation from both rest and task conditions, spanning spectral, aperiodic, and nonlinear domains.

#### 5.1.1 Broadband Spectral Features

Meditation was characterised by elevated theta–alpha power relative to both rest and task. Discrete theta and alpha bands uniquely distinguished the three states: theta activity was higher during meditation compared to rest, whereas alpha power was higher during meditation compared to the task. Interestingly, meditation resembled rest in alpha and task in theta, yielding the theta–alpha ratio as a distinguishing composite marker.

Enhanced theta likely reflects sustained internal concentration and executive regulation (Magosso et al., 2021; Nair et al., 2017), consistent with both concentrative meditation (Aftanas & Golocheikine, 2001; DeLosAngeles et al., 2016) and working memory literature (Cona et al., 2020; Roberts et al., 2013; Sauseng et al., 2010). Similar alpha levels between meditation and rest may reflect trait-level internalisation in long-term practitioners (Cahn et al., 2010; Cahn & Polich, 2006). Together, increased theta–alpha power suggests internally focused (Kakumanu et al., 2018; Sharma et al., 2023) with enhanced internal–external attention regulation (Magosso et al., 2021), consistent with a stable, relaxed yet cognitively engaged state (Aftanas & Golocheikine, 2001; Benedek et al., 2014; Lomas et al., 2015).

#### 5.1.2 Aperiodic Features

Aperiodic parameters, particularly IRASA intercept, differentiated meditation from both rest and task conditions. Meditation showed stronger divergence from rest than from task in aperiodic measures, suggesting modulation of broadband neural excitability beyond oscillatory changes.

Discrepancies between IRASA and FOOOF-derived slope/intercept estimates likely reflect methodological differences in separating periodic and aperiodic components (Donoghue et al., 2020; Wen & Liu, 2016). IRASA directly isolates fractal activity via resampling, whereas FOOOF models aperiodic components within a parametric fit, potentially yielding divergent estimates. (Gerster et al., 2022). The relative scarcity of aperiodic analyses in meditation research underscores the importance of including such measures to characterise state-level dynamics.

#### 5.1.3 Nonlinear Measures

Nonlinear measures robustly differentiated meditation from both rest and task states. Elevated DFA during meditation indicates increased long-range temporal correlations (Linkenkaer-Hansen et al., 2001), reflecting persistent and temporally structured neural dynamics associated with a better-synthesised self-related identity (Sugimura et al., 2021). In contrast, reduced DFA during the task likely reflects flexible, adaptive processing required for externally oriented demands (Hirekhan et al., 2019).

Meditation was associated with reduced Higuchi fractal dimension and permutation entropy relative to both comparison states. While reduced complexity is often associated with diminished consciousness, in this context, it likely reflects increased neural regularity and a more structured internal focus rather than reduced awareness. The pattern suggests the suppression of irrelevant network fluctuations and a reduction in neural noise to stabilise internal attention (Aftanas & Golocheikine, 2002).

Contradictory findings in prior complexity studies, with both reduced (Aftanas & Golocheikine, 2002; Vyšata et al., 2014; Young et al., 2021) and enhanced complexity (Kakumanu et al., 2018; Walter & Hinterberger, 2022) may stem from practice heterogeneity, frequency-specific computations (Irrmischer et al., 2018), and unaccounted expertise differences. Kakumanu et al. highlighted the nuances of proficiency-based state-trait differences across two groups of advanced meditators: The teachers group showed increased complexity from baseline, while the Senior meditators group showed no change. They also found an inverse relationship between the duration of meditation experience and changes in complexity (Kakumanu et al., 2018). Our cohort comprised a mix of experienced meditators, including a few teachers, with all having more than 8 years of experience.

Collectively, the neurodynamic core identified here reflects an internally dominant brain state marked by elevated theta-alpha power, increased aperiodic power (higher Intercept IRASA), enhanced temporal persistence (higher DFA), reduced complexity and entropy (lower Higuchi and Permutation Entropy). This configuration suggests a highly structured, temporally coherent, and internally stabilised neural state distinct from both passive rest and active task engagement.

### 5.2 Stability of meditation-specific neural correlates

Reliability is essential for validating meditation-specific neural markers. Our findings demonstrate strong stability across multiple potential sources of variability.

#### 5.2.1 Intra-subject stability

Previous studies have indicated strong temporal consistency of EEG linear and non-linear features during resting states (Ding et al., 2022; Duan et al., 2021; Suárez-Revelo et al., 2015), mental conditions (Li et al., 2024; Uudeberg et al., 2025), and cognitive tasks (McEvoy et al., 2000). We observed high inter-session reliability of the meditation-specific EEG features across days, particularly during the Actual M session, both during silent and guided meditation periods. Practice M session showed moderate to good reliability (see Supplementary file, Figure 3), suggesting stabilisation of the meditative state with repetition and reduced acclimatisation effects.

#### 5.2.2 Inter-subject stability

Individual differences based on personality traits (Buric et al., 2022), data quality and denoising methods (Lopez et al., 2023), state-trait effects (Treves et al., 2024) are known to influence neural and behavioural outcomes (R. Tang & Braver, 2020). High inter-subject correlation and reduced dispersion indicate convergence of multivariate neural profiles across individuals. Increased alignment on day two suggests progressive stabilisation of the induced meditative state. Even the Practice M session showed high inter-subject correlation and reduced dispersion with no difference across two days (Supplementary File, Figures 4.1 and 4.2)

#### 5.2.3 Meditative states

Though most studies use a guided approach, which has been shown to be beneficial in novice or short-term meditators (J. Han et al., 2025; Jo et al., 2024). The differential effect of guided versus unguided or silent meditation has not been studied. No significant differences emerged between guided and silent meditation in experienced practitioners for both Actual M and Practice M sessions (Supplementary file, Figure 5). This indicates that the identified neural core reflects state-level dynamics independent of instruction-dependent effects, an expected outcome in long-term meditators. However, this is an important reliability consideration when studying short-term or novice meditators.

#### 5.2.4 Practitioner’s expertise

Experience-based neural differences are widely reported in the literature, highlighting neuroplasticity mediated by hours of practice (Baron Short et al., 2010; Brefczynski-Lewis et al., 2007; Corby et al., 1978). Despite classifying practitioners by total practice hours, no reliable differences emerged between advanced and intermediate groups on either day during the guided and silent meditation periods for both Actual M and Practice M sessions (Supplementary file, Figure 6). This suggests that once a threshold of proficiency is reached, the meditative neurodynamic profile stabilises and becomes independent of incremental experience.

### 5.3 A Reliability-Informed Experimental Framework

Most meditation studies rely on single-session designs and passive baselines, or on a control group (novices or practitioners at different levels of experience). By employing repeated sessions across two days, including both passive and active comparators, and restricting the sample to experienced practitioners, the present design isolates state-dependent neural signatures while explicitly quantifying reliability. The inclusion of both rest and task comparators proved critical: spectral features more strongly differentiated meditation from task, whereas aperiodic components distinguished meditation from rest more robustly. These findings underscore the importance of examining multiple neurodynamic domains to comprehensively characterise meditation.

Repeating meditation sessions within and across days facilitated the assessment of intra-and inter-subject reliability, accounting for day-to-day differences. To ensure they entered a meditative state despite the controlled laboratory setting and to maintain uniformity across subjects with varying levels of expertise, we provided them with tradition-specific guided meditation, followed by silent meditation. Notably, the high intra- and inter-subject stability of meditation-specific EEG features, with no difference between guided and silent meditation periods, even across different levels of expertise, objectively verifies that the meditators are experienced and indicates that the identified neural correlates are specific to the meditative state.

## 6 Strengths, Limitations and Future Implications

The study provides a robust experimental framework for examining valid and reliable EEG-based correlates of meditation distinct from both an active control (a cognitive task) and a passive control (resting state) within a specific meditation tradition. A major strength of this study is the systematic evaluation of reliability across sessions, individuals, meditative formats, and expertise levels.

Further, this study involving long-term, proficient BKRY meditators provides guidelines for future EEG research to identify and address potential factors that underlie inconsistent findings in contexts such as meditation. To control for confounding factors such as acclimatisation and expectancy, it is recommended to include a practice or dummy session before the actual measurement.

We identified EEG features that differentiate meditative states from both resting and task states; the choice of EEG features determines how well they capture these differences. For example, spectral features distinguish meditation from task more effectively than from rest, whereas aperiodic components distinguish meditation better from rest than from task, showing more pronounced and widespread differences. This is due to differential properties reflected by distinct EEG features. Oscillatory power reflects rhythmic synchronisation associated with various cognitive, behavioural, and physiological processes (Buzsáki, 2006), whereas aperiodic activity captures the broadband excitation–inhibition balance (Donoghue et al., 2020; Gao et al., 2017) and nonlinear measures reflect scale-free dynamics related to irregularity, adaptability, and self-organisation (Klonowski, 2009). This highlights the importance of including a neurodynamic spectrum encompassing power spectral densities, aperiodic and nonlinear measures to obtain a comprehensive view of the brain’s internal neurodynamic state. We observed significantly higher power in the theta-alpha band during meditation than during rest or task, and lower power in the low-beta band relative to rest, and in the high-beta band relative to task. This highlights that overlapping regions across bands are essential for studying complex phenomena such as meditation. Also, to establish the robustness of the measures, intra- and inter-individual variability should be examined.

Despite the study’s strengths, there remains scope for further improvement. Inclusion of a meditation-naïve comparator group and phenomenological measures would further strengthen interpretability. The study could have employed a longer design rather than only consecutive days. Future work should integrate subjective experience with objective neurodynamic markers and extend this framework over a longer period of time across diverse contemplative traditions.

## Conclusion

This study identifies a reproducible neurodynamic core of meditation distinct from rest and cognitive task states, that remains stable across methodological and individual variability. In proficient practitioners, meditation is characterised by internally oriented, temporally persistent, regular, less complex and structured neural dynamics spanning spectral, aperiodic, and nonlinear domains. The importance of such neuronal organisation and structure for consciousness of the meditative state aligns well with the assumed role of form or organisation as a distinct dimension of consciousness, alongside content and level/state, as postulated by the Temporo-spatial theory of consciousness (TTC) (Northoff & Huang, 2017; Northoff & Zilio, 2022). The findings advance the construct validity and measurement reliability of the EEG-based markers of meditative state and provide a methodological template for future contemplative neuroscience research.

## Disclosures & Declarations

### Ethical Approval

Ethical approval was obtained from the Ethics Committee of NIMHANS Institute Human Research (Ethics Approval Number: NIMH/DO/Ethics sub-committee (BS) meeting/2024) and S-VYASA Institute (Ethics Approval Number: RES/IEC-SVYASA/337/2024. Written informed consent was obtained from all participants before participation.

## Consent from participants

Written informed consent was obtained from all participants before participation.

## Conflict of interest

The authors have no relevant financial or non-financial interests to disclose.

## Funding

No funding was received to assist with the preparation of this manuscript.

## Acknowledgement

The authors would like to acknowledge Dr Amruth Sagar for assistance with the initial setup and a few data acquisitions.

## Data availability

Currently, data is not publicly available. Will be made available on request.

## CTRI Registration

CTRI/2024/04/065384

## Notes

### Competing Interest Statement

The authors have declared no competing interest.

### Summary of Updates

The revised submission is just an update with the author and their affiliations.

