## Supplementary file for "The Neurodynamic Core of Meditation: Dissociating Meditation from Rest and Task in a Reliability-based EEG study"

1. Title of the article: **The Neurodynamic Core of Meditation: Dissociating Meditation from Rest and Task in a Reliability-based EEG study**
2. Name by which each contributor is known with highest qualifications and institutional affiliations:
   a. **Praerna Chowdhury,** Scientist-C, Centre for Consciousness Studies, Department of Neurophysiology, NIMHANS, Bangalore-560029, INDIA.
   b. **Ramajayam Govindaraj,** Assistant Professor, Indian Knowledge system & Mental Health Applications Centre (IKSMHA), Indian Institute of Technology (IIT), Mandi-175075, Himachal Pradesh, INDIA.
   c. **Arun Sasidharan,** Scientist-D, Centre for Consciousness Studies, Department of Neurophysiology, NIMHANS, Bangalore-560029, INDIA.
   d. **Apar A Saoji,** Principal – The School of Yoga and Naturopathic Medicine, Swami Vivekananda Yoga Anusandhana Samsthana, Bangalore-560019.
   e. **Ravindra PN,** Additional Professor, Centre for Consciousness Studies, Department of Neurophysiology, NIMHANS, Bangalore-560029, INDIA.
   f. **Georg Northoff,** The Royal’s Institute of Mental Health Research & University of Ottawa, Brain and Mind Research Institute, Centre for Neural Dynamics, Faculty of Medicine, University of Ottawa, 145 Carling Avenue, Rm. 6435, Ottawa, ON K1Z 7K4, Canada
   g. **Bindu M Kutty,** Senior Professor, Centre for Consciousness Studies, Department of Neurophysiology, NIMHANS, Bangalore-560029, INDIA.
3. The name, address, phone numbers, facsimile numbers and e-mail address of the contributor responsible for correspondence about the manuscript;

**Dr. Ramajayam Govindaraj,**

**Mob: 07760525398**

**Address:** Indian Knowledge system & Mental Health Applications Centre (IKSMHA), Indian Institute of Technology (IIT), Mandi-175075, Himachal Pradesh, INDIA.

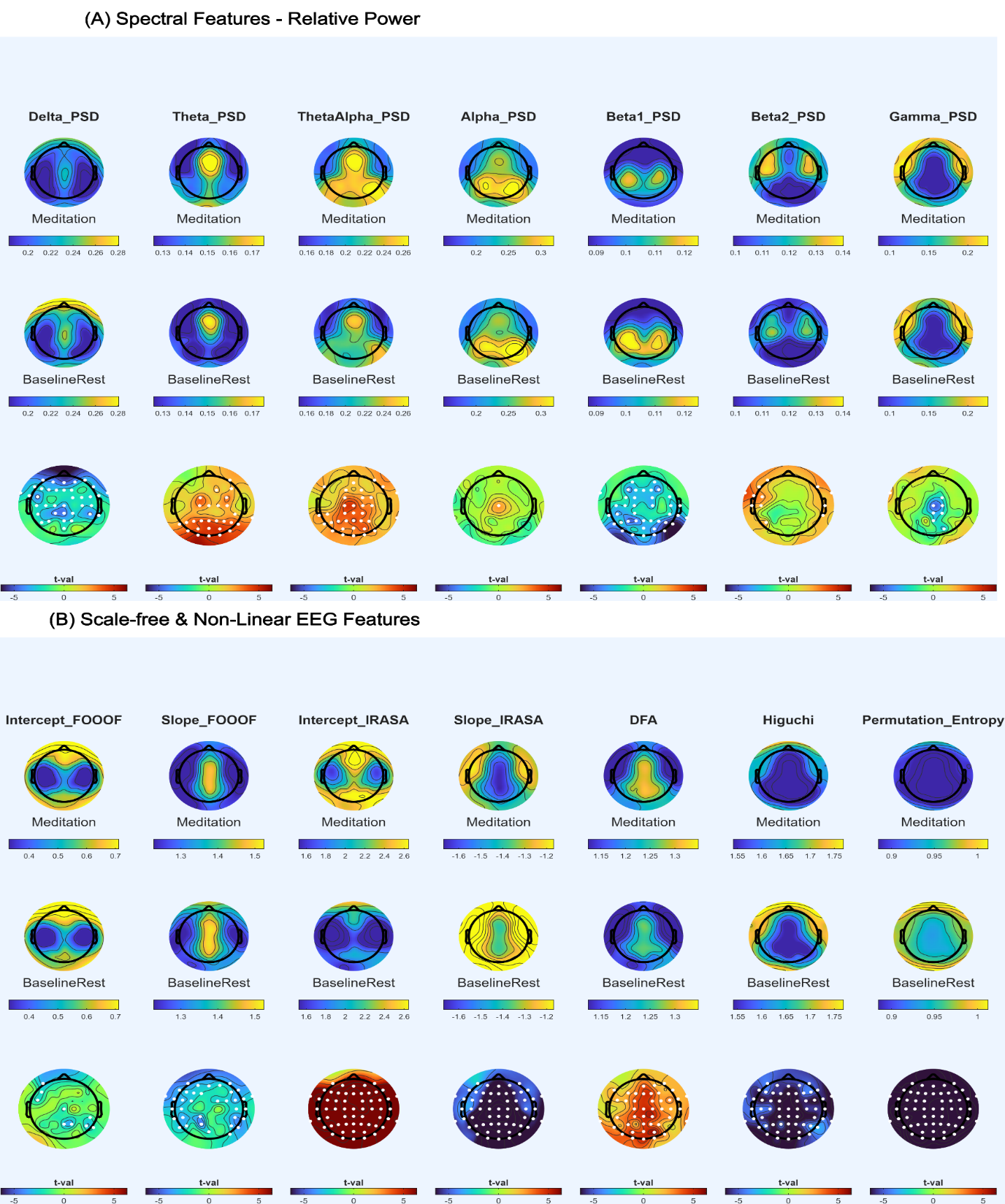

**Fig. 1** Topo-based comparison between Meditation and Baseline Rest across Broadband Spectral, Aperiodic, Complexity and Information-related EEG Features

Meditation – all Meditation sessions, including guided and silent meditation, averaged at each electrode level; Baseline Rest – eyes open baseline rest on two days, averaged at each electrode level. The top two rows present the average power spectra for each condition, while the bottom row shows the statistical difference between the two conditions as t-values, and white dots indicate electrode locations with statistically significant differences. A. Spectral Features and B. Scale-free and Non-linear EEG Features

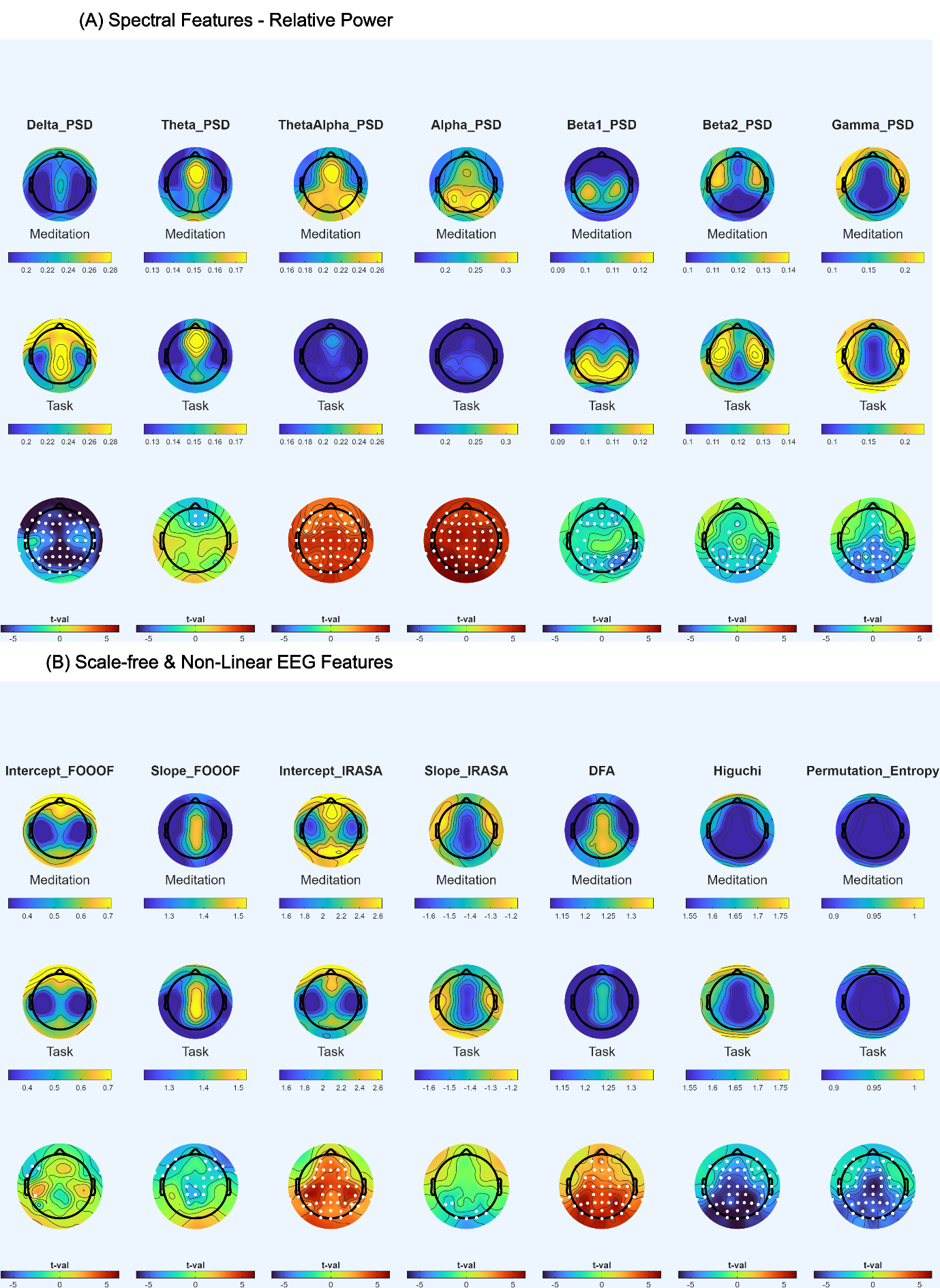

**Fig. 2** Topo-based comparison between Meditation and Cognitive Task across Broadband spectral, Aperiodic, Complexity and Information-related EEG Features

Meditation – all Meditation sessions, including guided and silent meditation, averaged at each electrode level; Task – working memory task on two days, averaged at each electrode level. The top two rows present the average power spectra for each condition, while the bottom row shows the statistical difference between the two conditions as t-values, and white dots indicate electrode locations with statistically significant differences. A. Spectral Features and B. Scale-free and Non-linear EEG Features

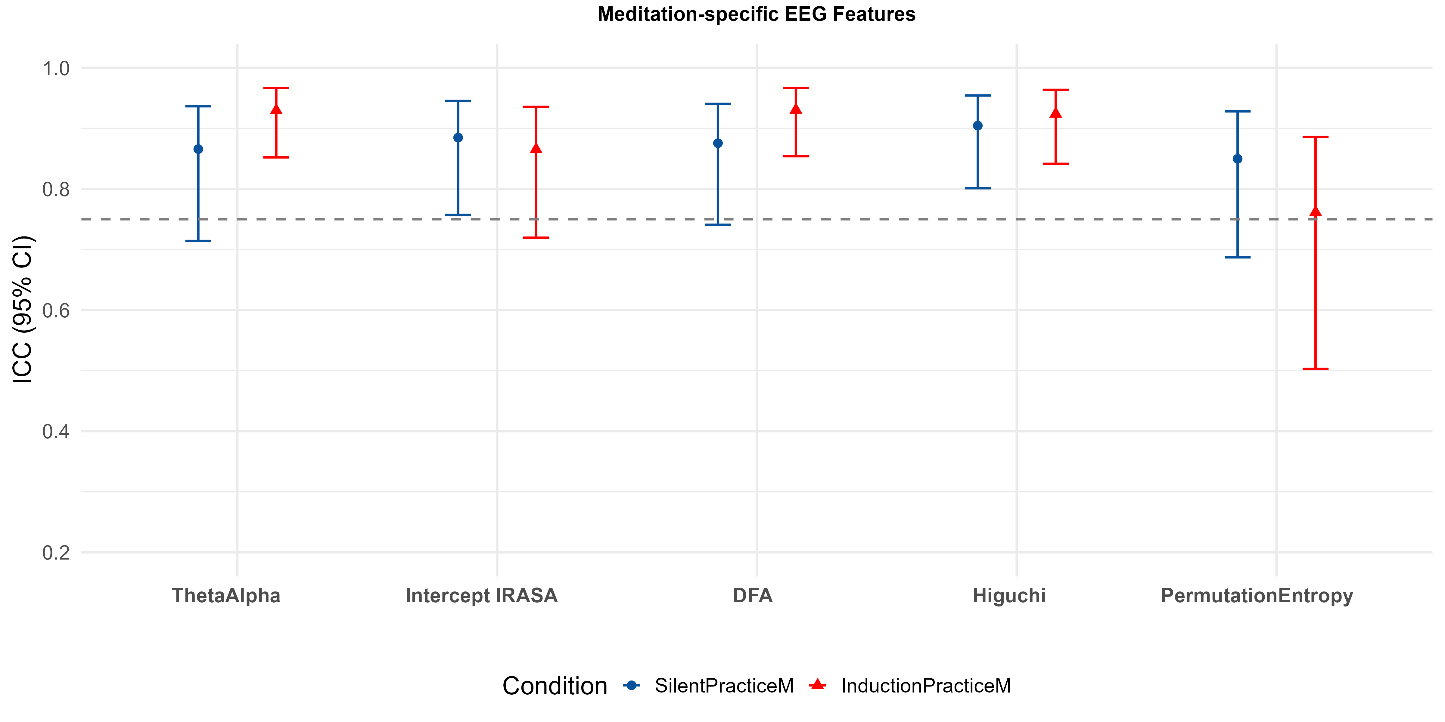

**Fig. 3** Intra-subject stability of meditation-specific EEG features during the silent and guided meditation periods of Practice M session across days

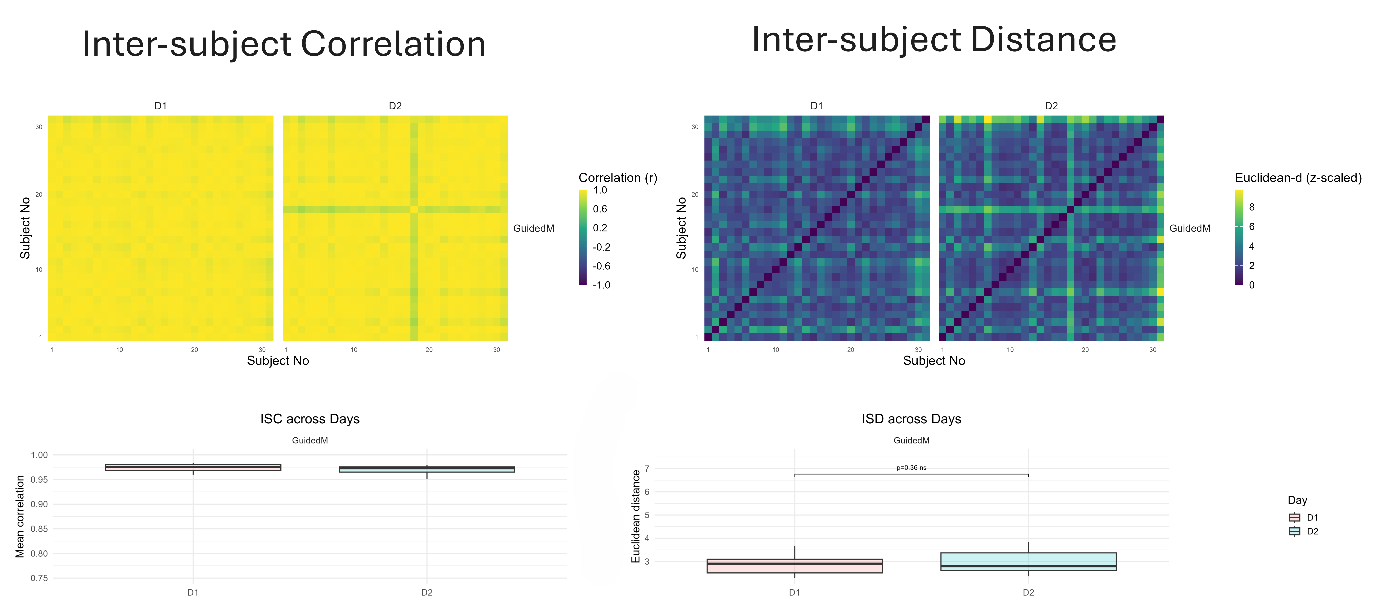

**Fig. 4.1** Inter-subject stability of meditation-specific EEG features during the guided meditation period of the Practice M session across days

Top Row: Inter-subject correlation (ISC, left) and Inter-subject distance (ISD, right) matrices computed from multivariate EEG features for day 1 (D1) and day 2 (D2). Each cell represents pairwise similarity or distance between subjects. Bottom row: Subject-wise means ISC and ISD across days. Boxplots show median and interquartile range; brackets indicate paired comparisons across days with FDR-corrected *p*-values.

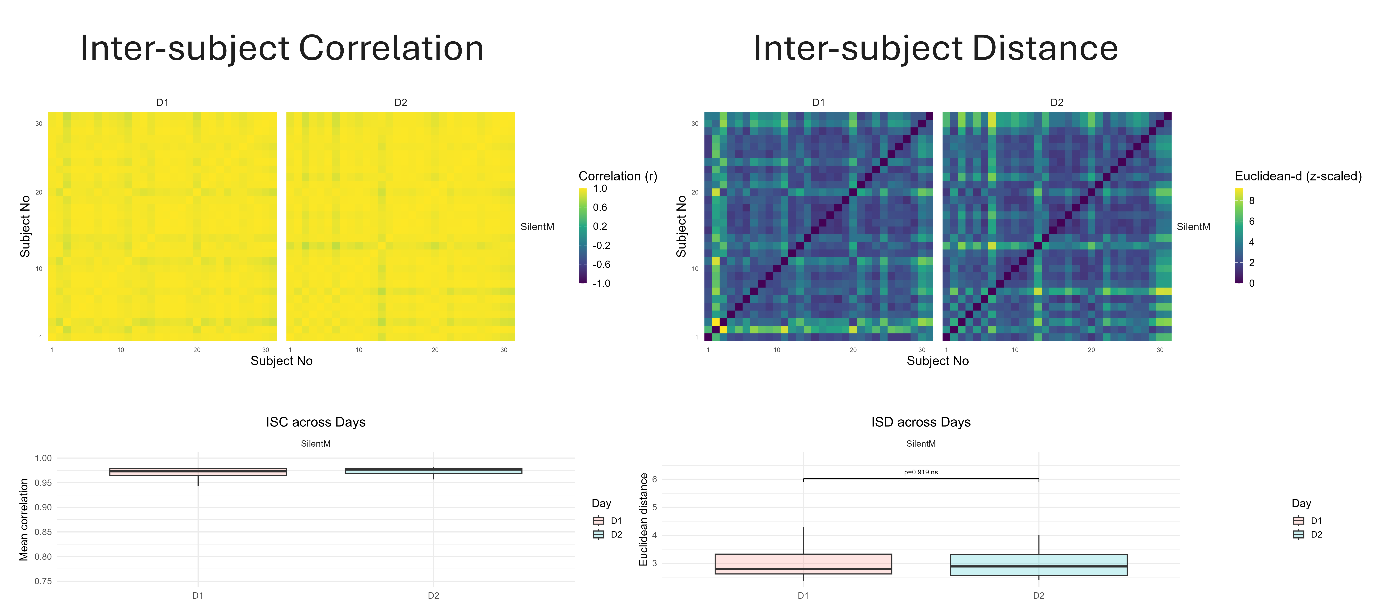

**Fig. 4.2** Inter-subject stability of meditation-specific EEG features during the silent meditation period of the Practice M session across days

Top Row: Inter-subject correlation (ISC, left) and Inter-subject distance (ISD, right) matrices computed from multivariate EEG features for day 1 (D1) and day 2 (D2). Each cell represents pairwise similarity or distance between subjects. Bottom row: Subject-wise means ISC and ISD across days. Boxplots show median and interquartile range; brackets indicate paired comparisons across days with FDR-corrected *p*-values.

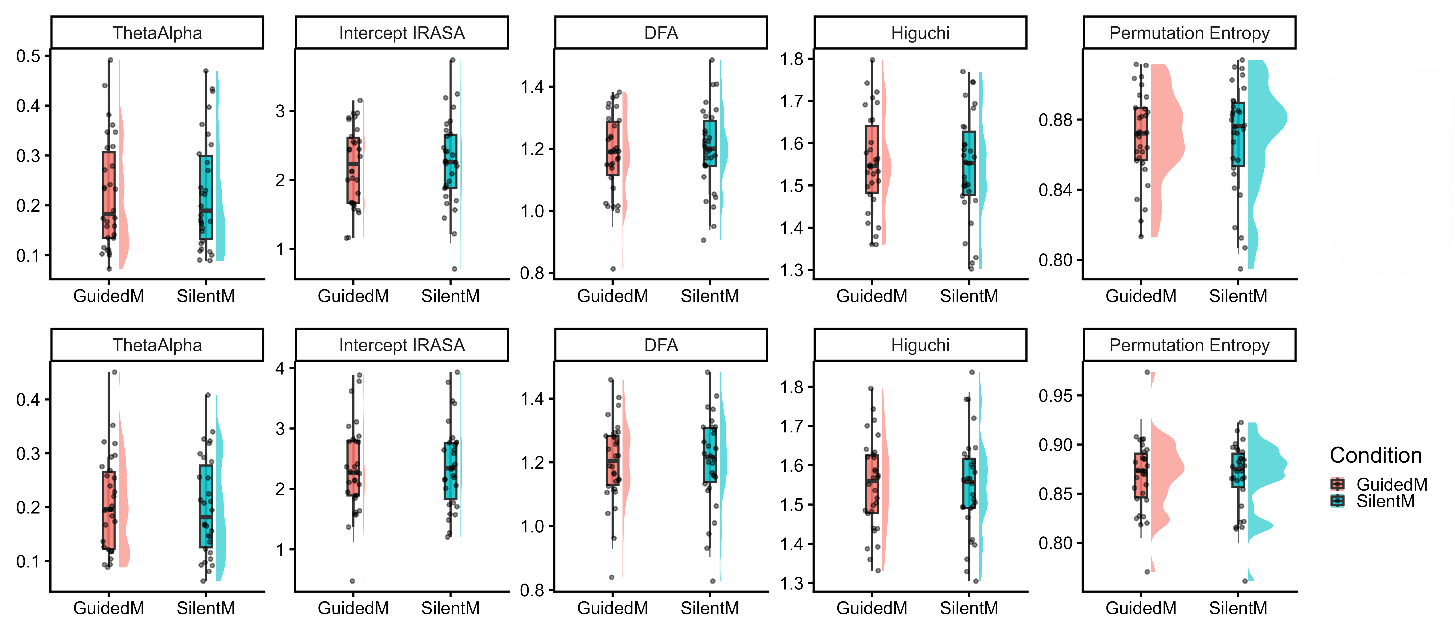

**Fig. 5** Raincloud plots with Yuen’s robust t-test (20% trimmed means) with FDR correction to assess differences between Guided and Silent periods of the Practice M session

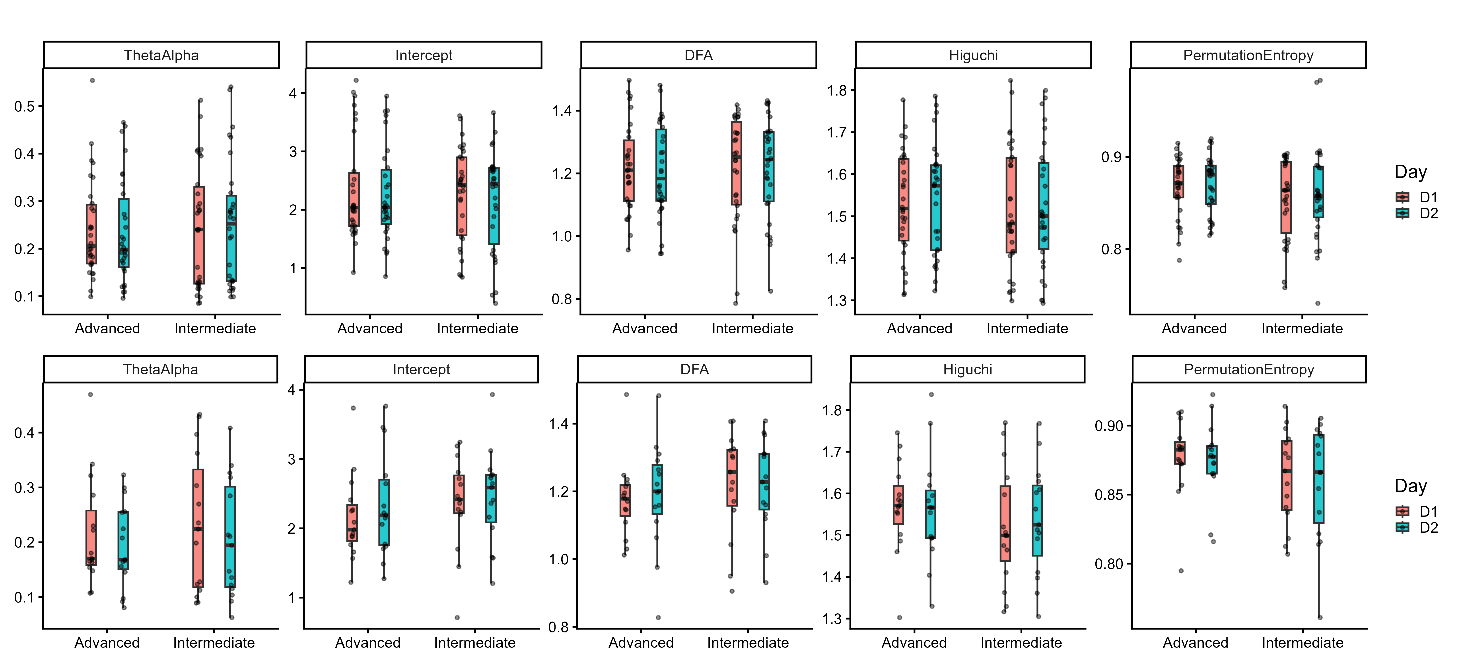

**Fig. 6** Raincloud plots with Yuen’s robust t-test (20% trimmed means) with FDR correction to assess differences between Advanced and Intermediate meditators, during guided and silent periods of Practice M session
